# How the Azadithiolate Ligand Impacts O_2_-Stability of Group B [FeFe]-Hydrogenase ToHydA

**DOI:** 10.64898/2026.04.16.719040

**Authors:** Subhasri Ghosh, Chandan K. Das, Shuvankar Naskar, Lars V. Schäfer, Thomas Happe

## Abstract

[FeFe]-hydrogenases are metalloenzymes that catalyze the reversible oxidation and production of H_2_, making them potential candidates for sustainable energy solutions. However, their practical application is restricted by their extreme O_2_ sensitivity, which leads to irreversible active site degradation. A newly characterized Group B hydrogenase, ToHydA from *Thermosediminibacter oceani*, has exhibited exceptional O_2_-stability even after long-term exposure to air. In ToHydA, the highly conserved proton-transporting cysteine (C212) safeguards the H-cluster from O_2_-induced degradation by formation of the H_inact_ state. In this study, we investigate the effects of replacing the azadithiolate (ADT) ligand of the [2Fe]_H_ subcluster by propanedithiolate (PDT), revealing that this substitution prevents the formation of the H_inact_ and H_trans_ states observed in wild-type ToHydA (bearing the ADT ligand). By combining ATR-FTIR spectroscopy and molecular dynamics (MD) simulations, we show that a favorable interaction between the nitrogen bridgehead of the ADT ligand and the C212 sidechain is crucial for stabilizing these states. The absence of this interaction in ToHydA^PDT^ (bearing the PDT ligand) prevents the C212 sidechain from approaching the Fe_d_ center of [2Fe]_H_, thereby reducing H_inact_ accumulation. Moreover, as-isolated ToHydA^PDT^ predominantly exhibits the H_hyd_ state, which is unusual for [FeFe]-hydrogenases with bound PDT ligand. These findings demonstrate how ligand substitution at the [2Fe]_H_ site of ToHydA affects the structural dynamics, offering detailed molecular insights into the ligand-dependent modulation of [FeFe]-hydrogenases.

## Introduction

Hydrogenases are metalloenzymes that catalyze both the production and oxidation of molecular hydrogen.^1,2^ Based on the metal composition of their active sites, these enzymes are classified into [Fe]-hydrogenases, [NiFe]-hydrogenases, and [FeFe]-hydrogenases, with the latter being the most efficient in catalyzing H_2_ production.^2–4^ The active site of [FeFe]-hydrogenases, known as the H-cluster, consists of a canonical [4Fe4S] cluster ([4Fe]_H_) and a unique diiron cluster ([2Fe]_H_).^5–9^ The [4Fe]_H_ cluster is coordinated by four cysteine residues, one of which also serves to link it to the [2Fe]_H_ diiron cluster. Each iron atom in the [2Fe]_H_ cluster is coordinated by a carbon monoxide (CO) and a cyanide (CN^-^) ligand and a third CO ligand is found in a bridging position between both Fe ions. An additional azadithiolate bridge links the Fe ions, the amine group serving as a proton relay. The iron atoms are designated as proximal Fe (Fe_p_) and distal Fe (Fe_d_) based on their positions relative to the [4Fe]_H_-cluster. In the resting state, known as the H_ox_ state, the H-cluster features a combination of Fe_d_(I) and Fe_p_(II) oxidation states, with the Fe_d_ center possessing an open coordination site that serves as the ligand-binding site during catalytic turnover.^2,10^

The catalytic states of [FeFe]-hydrogenases have been widely studied. Throughout the catalytic cycle, the oxidation states of the two iron atoms in the [2Fe]_H_ cluster switch between Fe(I) and Fe(II), in conjunction with the 1+/2+ redox transitions of the [4Fe]_H_ cluster. Through successive reduction and protonation of the H_ox_ state, the system transitions to the H_red_ and H_sred_ states, characterized by a [Fe_p_(I)-Fe_d_(I)] configuration.^2^ However, the precise structure and degree of protonation in these states remain subjects of debate, with alternative designations as H_red_H^+^ and H_sred_H^+^, suggesting additional protonation at the ADT bridge.^11,12^ A structural rearrangement of the H_sred_H^+^ state results in the coordination of a terminal hydride at the Fe_d_ site, giving rise to the H_hyd_ state with a [Fe_p_(II)-Fe_d_(II)] configuration which is an essential intermediate in the catalytic cycle of [FeFe]-hydrogenases.^13–19^

The Fe_d_ site of [FeFe]-hydrogenases can bind various ligands, such as CO, CN^-^, SH^-^, and O_2_, which can lead to either reversible inactivation or irreversible degradation.^20–28^ In particular, degradation due to O_2_ exposure presents a significant challenge for large-scale biotechnological applications of these enzymes.^29^ Although most [FeFe]-hydrogenases are highly sensitive to O_2_, a few exceptions have been identified: CbA5H from *Clostridium beijerinckii*, CpIII from *Clostridium pasteurianum* and ToHydA from *Thermosediminibacter oceani* are recognized as O_2_-tolerant [FeFe]-hydrogenases.^30–33^ In these enzymes, under oxidative conditions, the thiol group of a protontransporting cysteine residue shifts to the Fe_d_ site, forming the H_inact_ state, which protects the active site from the damage caused by O_2_. While CbA5H belongs to the well-characterized Group A [FeFe]-hydrogenases, CpIII and ToHydA are Group B (M2a type) hydrogenases.^30,33,34^ ToHydA differs from Group A hydrogenases in its amino acid sequence, which results in distinct catalytic behavior and spectroscopic properties. In our recent study, we showed that, unlike CbA5H and the O_2_-sensitive Group A hydrogenases, ToHydA can form the H_inact_ state even in its resting state and stabilizes an intermediate, the H_trans_ state, during its transition to the active H_ox_ state. Both H_inact_ and H_trans_ share a [Fe_p_(II)-Fe_d_(II)] configuration (Figure 1A). We further identified the amino acid residues critical for H_inact_ formation, with particular emphasis on the differences between Group A and Group B (M2a) hydrogenases. From these findings, a tentative mechanism for H_inact_ formation has been proposed. Nonetheless, the common structural features of various groups of [FeFe]-hydrogenases, such as the proton-transporting cysteine and ADT ligand, could also influence this process, underscoring the need for a deeper understanding of the inactivation mechanism to guide the design of highly active, O_2_-stable biocatalysts for H_2_ production.^33^

**Figure 1.**
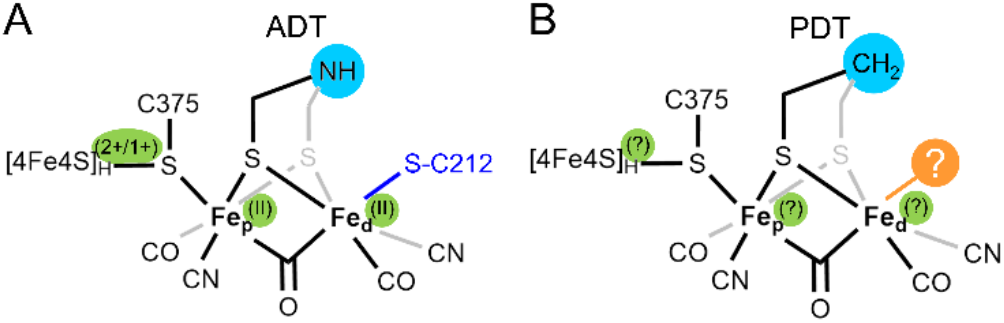
Schematic representations of A) the H_inact_/H_trans_ state, predominantly observed in the H-cluster of ToHydA^ADT^ (WT) under various conditions, and B) the H-cluster of ToHydA^PDT^ with unknown oxidation states and ligand at Fe_d_. C375 connects the [4Fe]_H_ and [2Fe]_H_ clusters, whereas C212 participates in proton transport and the formation of the H_inact_ and H_trans_ states.

To investigate the inactivation mechanism and catalytic features of ToHydA, this study focuses on the bridgehead position of the H-cluster by replacing the native ADT ligand with a propanedithiolate (PDT) ligand (Figure 1B). While the effects of this modification on catalytic properties have been studied in Group A hydrogenases, its impact in Group B hydrogenases has not yet been explored. We examined the different H-cluster states of the PDT variant using attenuated total reflection Fourier-transform infrared (ATR-FTIR) spectroscopy, which provides insights into the H_hyd_ state of ToHydA^PDT^ under conditions where this state is typically absent in Group A [FeFe]-hydrogenases. In addition, we carried out all-atom molecular dynamics (MD) simulations of both ToHydA^ADT^ and ToHydA^PDT^ variants to obtain an atomic-level understanding of the structural consequences of this modification, particularly during H_inact_ formation. Collectively, our findings provide deeper insight into the catalytic behavior of ToHydA, especially regarding H_inact_ formation, which could foster the future design of *de novo* [FeFe]-hydrogenases for biotechnological applications.

## Results and Discussion

The ToHydA apoprotein was heterologously expressed and produced in *E. coli*. Following purification, the protein was matured using synthetic diiron complexes, as previously described.^9,33^ In this study, two diiron complexes were utilized (see Figure 1): [Fe_2_[μ-(SCH_2_)_2_NH](CN)_2_(CO)_4_]^2-^, which contains the native ADT ligand and results in catalytically active wild-type ToHydA upon maturation, and [Fe_2_[μ- (SCH_2_)_2_CH_2_](CN)_2_(CO)_4_]^2-^, which incorporates the PDT ligand. The resulting proteins are referred to as ToHydA^ADT^ (identical to ToHydA WT) and ToHydA^PDT^, respectively. To investigate the formation of various H-cluster states in ToHydA^ADT^ and ToHydA^PDT^, ATR-FTIR spectroscopy was employed (Figure 2). In its as-isolated state (representing the semi-dried protein on the ATR crystal without any treatment), ToHydA^ADT^ predominantly exhibits the H_inact_ and H_trans_ states. Upon treatment with N_2_ and O_2_, it predominantly accumulates the H_inact_ state.^33^

**Figure 2.**
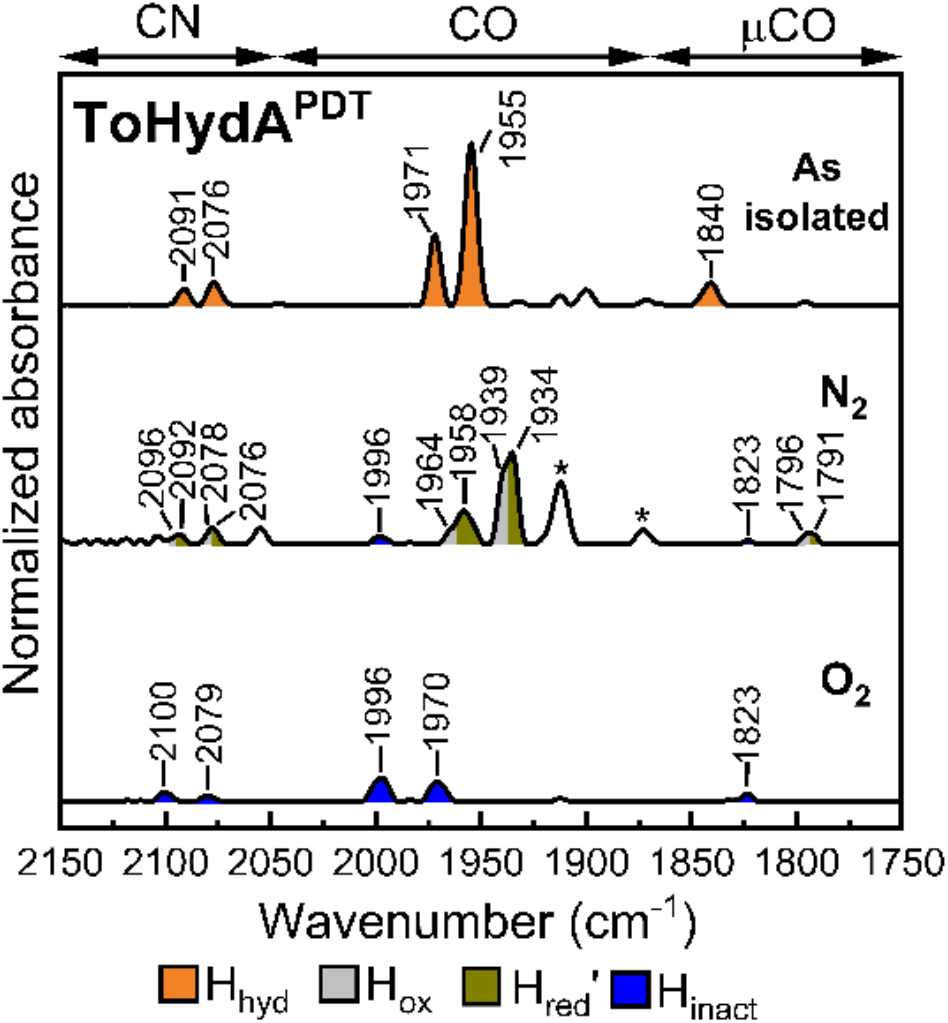
FTIR spectra of ToHydA^PDT^ in as-isolated state (upper panel), upon N_2_ (8 L/min) purging (middle panel) and after O_2_ (air, 2 L/min) purging (lower panel). The spectra of the proteins are normalized to the second amide band (1535 - 1545 cm^−1^). 0.4 - 0.5 mM protein samples were prepared in 100 mM Tris/HCl buffer at pH 8.0 containing 2 mM NaDT.

In contrast to ToHydA^ADT^, the as-isolated ToHydA^PDT^ accumulates a state that appears distinct from both the H_inact_ and H_trans_ states (Figure 2, upper panel). The FTIR spectrum of the asisolated ToHydA^PDT^ reveals a single μCO peak at 1840 cm^-1^, two CO peaks at 1955 cm^-1^ and 1971 cm^-1^, and two CN^-^ ligand peaks at 2076 and 2091 cm^-1^, indicating the presence of a single state. The terminal CO peaks of the newly observed state appear close to those of the hydride-bound H_hyd_, H_trans_ and H_ox_ states, indicating an H-cluster electronic configuration of either [4Fe]_H_^+^[Fe_p_(II)-Fe_d_(II)]_H_ (for H_hyd_ and H_trans_ states) or [4Fe]_H_^2+^[Fe_p_(I)-Fe_d_(II)]_H_ (for H_ox_ state).^10,17,35^ Moreover, the μCO peak of this state is blue-shifted relative to that of the H_ox_ state, suggesting ligand binding at the open coordination site.

In Group A hydrogenases, the H_hyd_ state is typically observed in catalytically inactive variants with disrupted proton transport pathways under reducing conditions.^17,19^ To elucidate the identity of the newly observed state, we generated a ToHydA C212A variant, in which the proton-shuttling cysteine (C212) was replaced by alanine, thereby disrupting the proton transfer pathway. The ToHydA C212A variant was matured with the diiron complex containing the ADT ligand, referred to here as C212A^ADT^, and characterized using ATR-FTIR spectroscopy. The spectrum of the as-isolated C212A^ADT^ variant suggests the accumulation of the H_ox_CO state primarily together with H_ox_ (Figure 3, upper panel); however, the new state described above was not observed under these conditions. Since the H_hyd_ state is known to form under reducing conditions,^13,33^ the C212A^ADT^ variant was treated with 25 mM NaDT, which led to the accumulation of the new state, supporting its assignment as the hydride-bound H_hyd_ state (Figure 3, lower panel).

**Figure 3.**
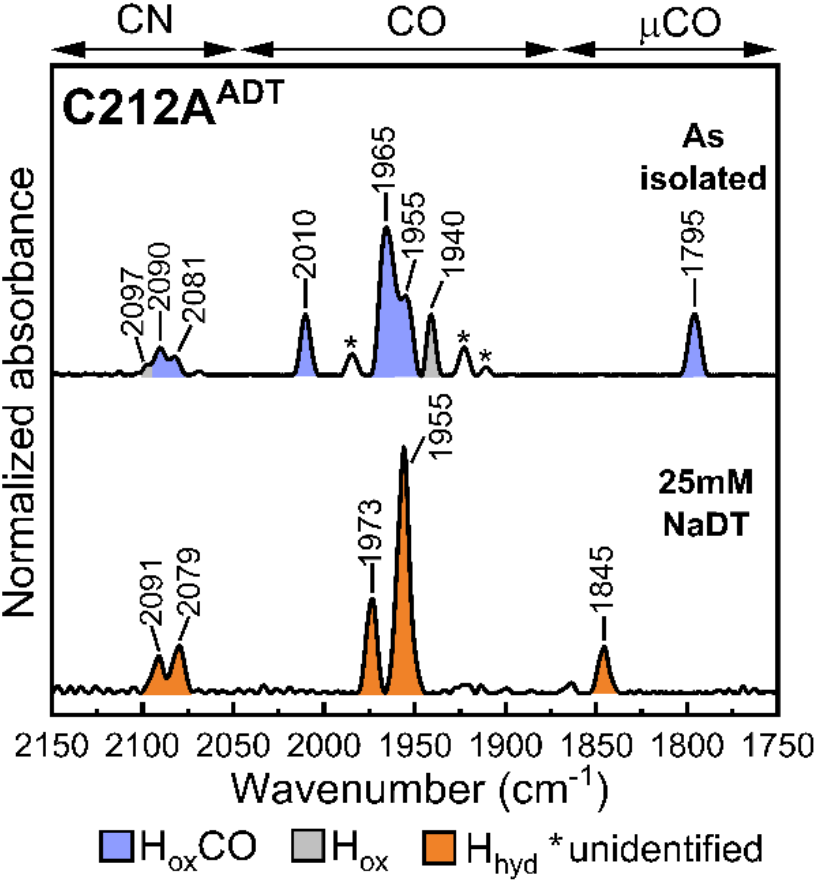
FTIR spectra of ToHydA C212A^ADT^ variant in as-isolated state (upper panel) and upon treatment with 25 mM NaDT (lower panel). The spectra of the proteins are normalized to the second amide band (1535-1545 cm^−1^). 0.4 - 0.5 mM protein sample was prepared in 100 mM Tris/HCl buffer at pH 8.0 containing 2 mM NaDT.

To further investigate the binding of a hydride ligand, we employed a primary isotope effect by exchanging H_2_O in the buffer with D_2_O. As the H_hyd_ state involves a terminal hydride, the isotope effect can be observed as a shift in the peak position of the bridging CO in the FTIR spectrum when D^-^ occupies the axial position at the Fe_d_ site instead of H^-^.^14,16,17,36^ In similar isotope exchange experiments, the µCO peak of ToHydA^ADT^ did not shift in the FTIR spectra, leading to the assignment of the H_trans_ state in ToHydA.^33^ The as-isolated ToHydA^PDT^ in D_2_O-containing buffer displayed the new state, as evidenced by similar peak positions of the terminal CO and CN^-^ ligands (Figure 4). However, the intensity of the bridging CO peak at 1840 cm^-1^ was significantly reduced, and a new, smaller peak appeared at 1831 cm^-1^ (Figure 4A, middle and right panel). This observation indicates that both H^-^ and D^-^ coordinate to the Fe_d_ in the protein preparation, although the overall intensity of the μCO peaks at 1840 and 1831 cm^-1^ is relatively low compared to the other CO and CN^-^ ligand peaks. A few unidentified peaks are observed in the as-isolated spectrum of ToHydA^PDT^ in D_2_O. These peaks are also present in the spectrum of the protein prepared in H_2_O and are distinct from the H_hyd_ state (SI Figure S1). Additionally, the FTIR spectrum in the range of 1000 to 4000 cm^-1^ showed peaks corresponding to both D_2_O and H_2_O (Figure 4A, left panel),^37^ suggesting that under the alkaline experimental conditions and in presence of H_2_ (2%), D_2_O is partially converted to H_2_O, as previously reported.^38^ This supports the notion that either H^-^ or D^-^ may coordinate to the Fe_d_. A similar isotope exchange experiment was performed for the C212A^ADT^ variant in the presence of 25 mM NaDT, prepared in D_2_O-containing buffer. In this variant, a peak at 1836 cm^-1^ was observed alongside the peak at 1845 cm^-1^, corresponding to the binding of D^-^ and H^-^, respectively (Figure 4B, middle and right panels). The presence of both H_2_O and D_2_O was again confirmed by their characteristic peaks in the range of 1000 to 4000 cm^-1^ (Figure 4B, left panel). These isotope exchange experiments strongly support that the new state is indeed the H_hyd_ state.

**Figure 4.**
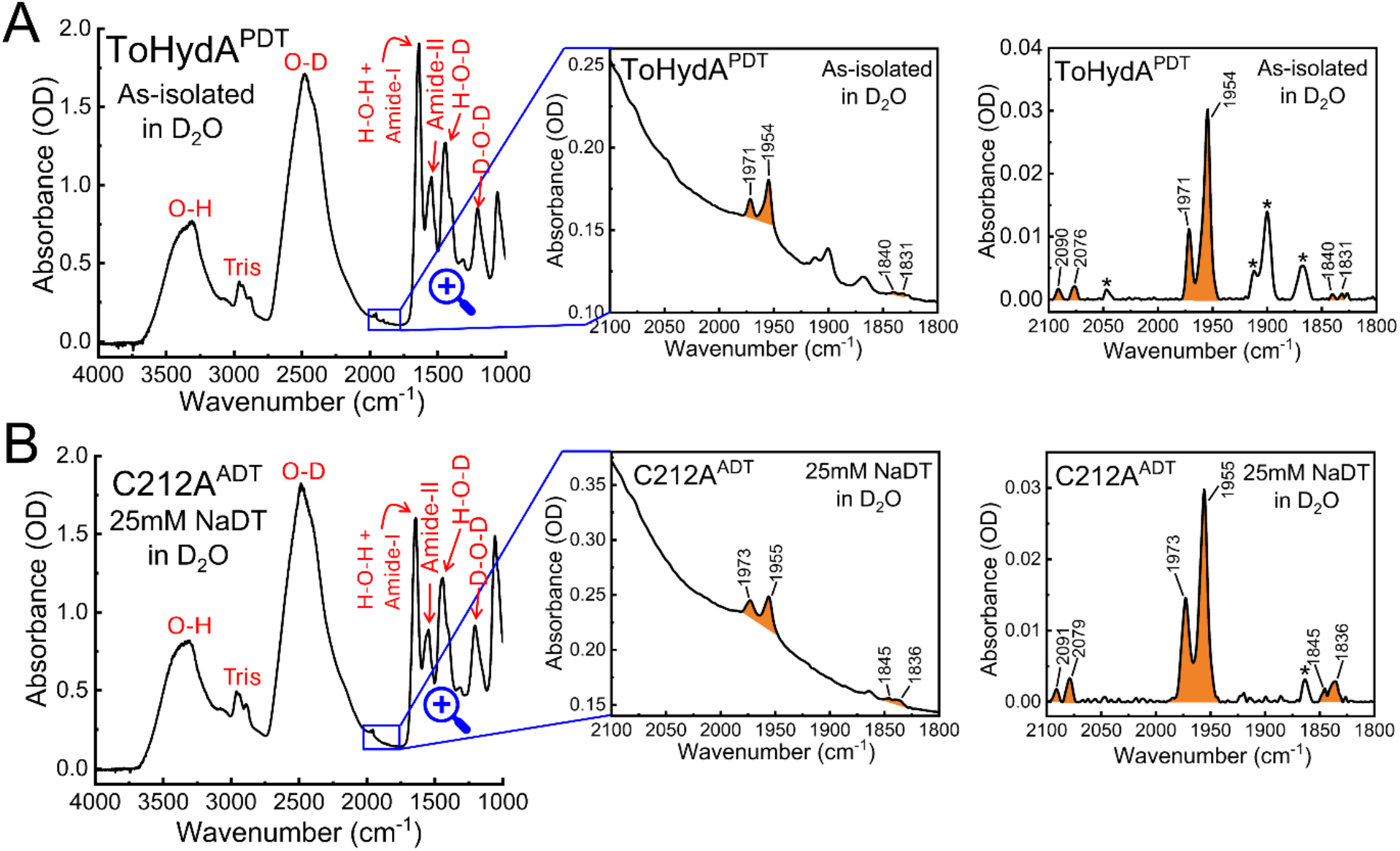
A) FTIR spectrum of ToHydA^PDT^ in the as-isolated state, B) FTIR spectrum of ToHydA C212A^ADT^ upon treatment with 25 mM NaDT prepared in 100 mM Tris/HCl buffer in D_2_O at pH 8.0. The left panel displays the full spectral range (4000–1000 cm^−1^), with peaks corresponding to the vibrational modes of H_2_O/D_2_O/HDO labeled in red. No post-processing (atmospheric compensation, baseline correction and normalization) was applied to the spectrum. The H-cluster region (2100-1800 cm^-1^) of the spectrum is shown in the middle panel. The right panel shows the baseline corrected H-cluster spectrum. 0.5 mM protein sample was prepared in 100 mM Tris/HCl buffer in D_2_O at pH 8.0 containing 2 mM NaDT. The H_hyd_ state peaks are colored orange.

We additionally characterized ToHydA^PDT^ by FTIR spectros-copy at pH 10. The as-isolated protein (prepared in 100 mM Tris/HCl at pH 10.0 containing 2 mM NaDT) exhibited the H_hyd_ state also at the higher pH (Figure S2). The H_hyd_ state was also retained in the presence of 25 mM NaDT at both pH 8 and pH 10 (Figure S2), further supporting that H_hyd_ is not converted to other reduced states even at elevated reducing pressure. Upon treatment with N_2_, FTIR spectroscopy showed that ToHydA^PDT^ primarily forms H_ox_-like states (Figure 2, middle panel). The broadening of peaks around 1934 cm^-1^ and 1958 cm^-1^ suggests the coexistence of either the H_ox_H or H_red_′ state alongside the H_ox_ state (together referred to as H_ox_-like states). However, unlike ToHydA^ADT^, the H_inact_ state is almost absent in N_2_-treated ToHydA^PDT^. Under H_2_ treatment, no H_hyd_ formation was observed at either pH 8 or pH 10 (Figure S2). Instead, the spectra at pH 8 displayed H_ox_-like states (similar to N_2_ treatment), and the H_red_′ state as the dominant species at pH 10. Thus, rather than promoting hydride formation, H_2_ shifts ToHydA^PDT^ toward the H_ox_-like and H_red_′ states, underscoring that the H_hyd_ state is favored under the reducing conditions of the as-isolated sample. Upon exposure to air, the H-cluster peaks are significantly diminished, indicating severe degradation of the H-cluster (Figure 2, lower panel). Only a small fraction of the protein (∼10%) is not degraded and forms the H_inact_ state (Figure S3).

As the thiol group of C212 participates in the H_inact_ formation of ToHydA to protect it from O_2_-induced degradation, understanding the structure and dynamics of the C212-bearing loop in To-HydA^ADT^ and ToHydA^PDT^ is crucial. To gain deeper atomic-level insights, molecular dynamics (MD) simulations were conducted using the AlphaFold-predicted model (ColabFold v1.3.0, AlphaFold2 using MMseqs2)^39,40^ of ToHydA^ADT^ (as described previously).^33^ The ToHydA^PDT^ variant was generated by replacing the ADT ligand in the WT structure with a PDT ligand. We analyzed a total of 1500 ns of MD trajectories for each system (see details in SI) and analyzed the distance between the sulfur atom of the C212 sidechain (C212:S_γ_) and the Fe_d_ atom in the [2Fe]_H_ cluster. In ToHydA^ADT^, the MD simulations show that the thiol group of C212 stays near the Fe_d_ site, whereas in ToHydA^PDT^, it is positioned further away (Figures 5A, B). The distance distributions also reveal differences in the dynamics of the C212 residue (Figure 5C; the most likely distances and the preferred orientation of C212 are depicted in Figures 5A and 5B). The simulations show that the presence of the ADT bridge is essential for the sulfur atom of C212 to closely approach Fe_d_ and form an H_trans_-like state. Close examination of the To-HydA^ADT^ simulations revealed that, unlike with PDT, a hydrogen bond can form between N_adt_ and C212:S_γ_, bringing the thiol group of C212 into close proximity of the Fe_d_ site (distance only 3 - 4 Å, see marked region in Figure 5C and representative configuration in Figure 5D).

**Figure 5.**
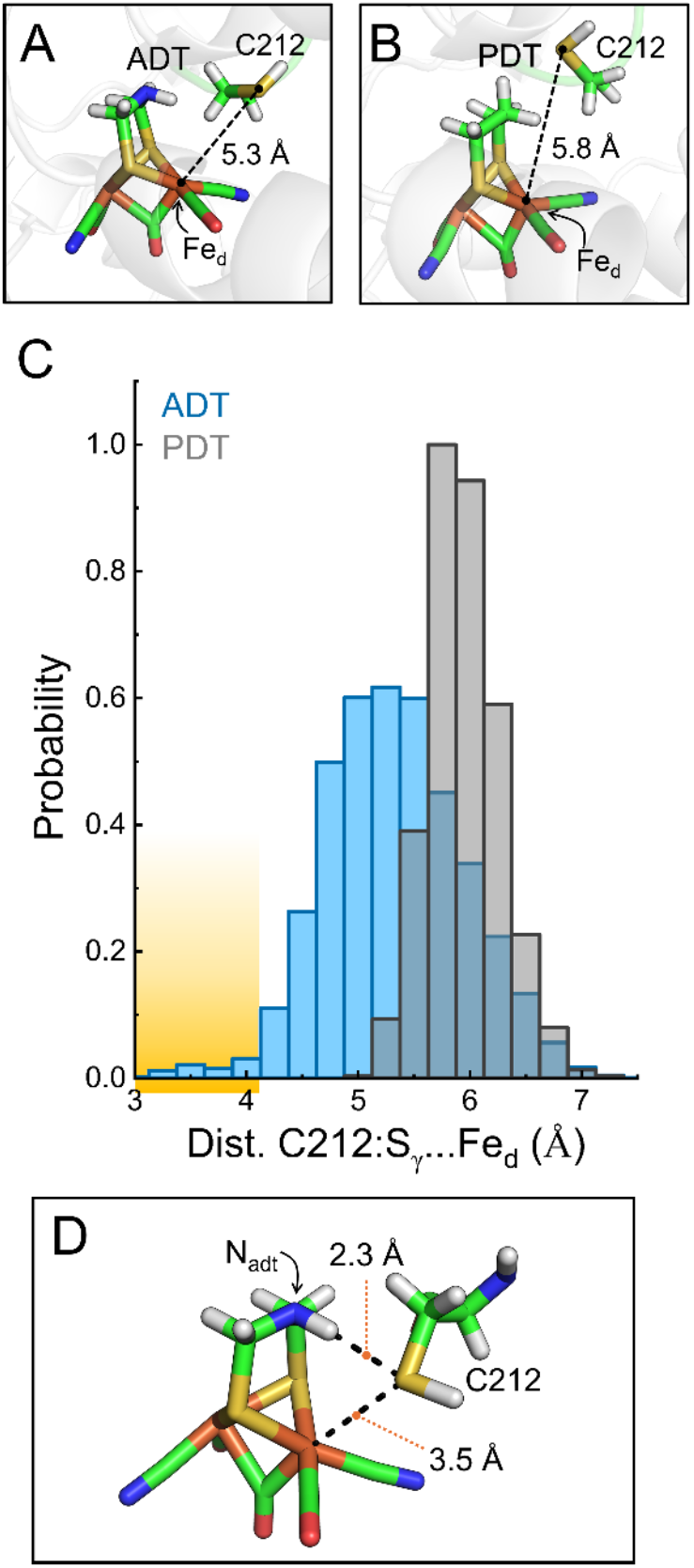
A) and B) Representative MD snapshots of ToHydA^ADT^ and ToHydA^PDT^, illustrating the most probable distance between C212:S_γ_ and Fe_d_. C) Distance distribution between C212:S_γ_ and Fe_d_ in ToHydA^ADT^ and ToHydA^PDT^. D) MD snapshot of ToHydA^ADT^ depicting an H_trans_-like state.

Although the population of H_trans_-like conformations is low in the MD simulation of ToHydA^ADT^, the observed accumulation of H_inact_ and H_trans_ states in ToHydA^ADT^ suggests a higher prevalence of conformations with short C212:S_γ_…Fe_d_ distances, stabilized by the interaction with the NH bridgehead. A comparable hydrogen bond interaction is also observed during inhibition through extrinsic sulfide binding.^26^ Taken together, the MD simulations support the notion that the formation of H_inact_ and H_trans_ in ToHydA^ADT^ is facilitated (Figure 6A). Similar H-bonding interactions between the ADT-amine and the second coordination sphere are known to be crucial for ligand binding and reactivity of the [2Fe]_H_.^41,42^

**Figure 6.**
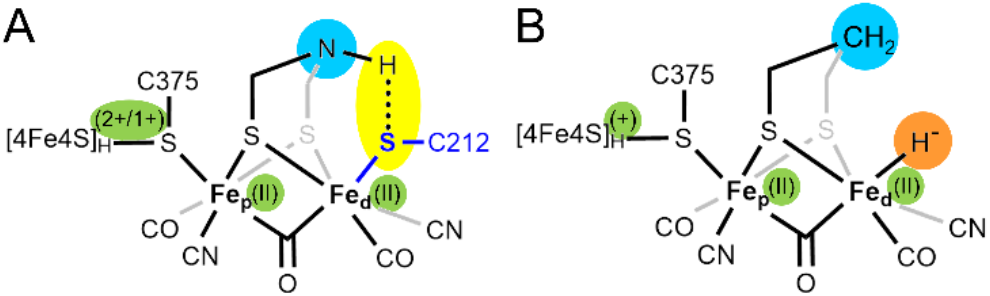
Schematic representation of A) the H_inact_/H_trans_ state of ToHydA^ADT^ (the dotted line shows the hydrogen bond between the bridgehead NH-group and C212:S_γ_), and B) the hydride-bound H_hyd_ state formed in ToHydA^PDT^.

The catalytic intermediate H_hyd_ typically appears in proton transport-disrupted variants of standard Group A [FeFe]-hydrogenases (such as CpI and CrHydA1), similar to the ToHydA C212A^ADT^ variant.^14,17,43^ In these variants, the lack of proton availability at the active site arrests catalytic turnover, resulting in accumulation of the H_hyd_ state. Although the PDT variant also disrupts proton transport, the absence of the H_hyd_ state in this variant remained unclear,^11,12^ until Depala *et al*.^44^ recently showed that the H_hyd_ state is formed in CrHydA1^PDT^ under significantly low potential (< −0.66 V). A more negative potential is required for the PDT variant to reduce the [2Fe]_H_ site, as the bridgehead CH_2_ group is less electronegative compared to the native NH bridgehead. Moreover, protonation of the amine bridgehead in the native ADT cofactor stabilizes the H_red_H^+^ and H_sred_H^+^ states followed by tautomerization to H_hyd_. In the PDT variant, this pathway is unavailable, as the methylene bridge-head cannot be protonated. Consequently, [2Fe]_H_ reduction in this variant must proceed via direct reduction to H_hyd_, requiring a substantially more negative potential. Nevertheless, in To-HydA^PDT^, the H_hyd_ state is formed under mild reducing conditions. While it is hard to predict the redox potential of [2Fe]_H_ in ToHydA compared to the Group A hydrogenases and why To-HydA^PDT^ forms H_hyd_ readily, it could be assumed that stability of the [Fe_p_(II)-Fe_d_(II)] electronic configuration (as shown in Figure 6B) of the [2Fe]_H_ cluster in ToHydA is one of the leading causes. Previously, in ToHydA^ADT^, states featuring ligand binding to Fe_d_, such as H_inact_, H_trans_, and the secondary inactive state, have demonstrated a preference for the [Fe_p_(II)-Fe_d_(II)] configuration.^33^ The presence of the H_hyd_ state in ToHydA^PDT^ further supports the idea of enhanced stability of the [Fe_p_(II)-Fe_d_(II)] electronic configuration, a prerequisite for the formation of the O_2_-stable H_inact_ state. In addition, the redox potentials of the auxiliary [4Fe4S] clusters may influence the redox states of the [2Fe]_H_ site.

While ToHydA and CpIII both belong to Group B [FeFe]-hydrogenases, they exhibit different H-cluster states under different conditions (such as N_2_, H_2_, O_2_ environment) and also differ in their inactivation properties.^32,33,45,46^ A direct one-to-one comparison is challenging due to the complex FTIR spectra of CpIII.^45^ In contrast to the spectral evidence observed for other enzymes, no fully reduced states, such as H_sred_H^+^ or H_hyd_ were identified. Consequently, while the H-cluster states of CpIII encompass the majority of the catalytic intermediates, it is either incapable of forming H_sred_H^+^ and H_hyd_, or these states are significantly destabilized in CpIII. In contrast, an H_trans_-like state has been identified, although the identity of the bound ligand remains unresolved. Additionally, the FTIR spectra of deuterated CpIII exhibit an overall very low intensity, with the μCO band at 1838 cm^-1^ being particularly diminished.^45^ Comparable low FTIR signal intensity has also previously been reported for other [FeFe]-hydrogenases under deuterium exchange conditions, complicating the identification of the respective states.^47^ In the case of ToHydA^PDT^, this effect is especially pronounced for the μCO peak of the H_hyd_ state (at 1840 cm^-1^). Nevertheless, FTIR spectroscopic characterization of the proton-transport disrupted C212A^ADT^ variant in presence of reducing agent supports the formation of the H_hyd_ state. However, further studies are required to elucidate the factors governing the formation of the [Fe_p_(II)–Fe_d_(II)] electronic configuration of the [2Fe]_H_ site in ToHydA.

Substitution of ADT with PDT is known to substantially impair, or even abolish, catalytic activity in [FeFe]-hydrogenases.^9^ Lachmann *et al*. recently reported that PDT variants of DdHydAB and CrHydA1 retain only a minor fraction of native H_2_-oxidation activity, with low but measurable current densities in protein film electrochemistry (PFE).^48^ Given the already very low H_2_-oxidation current density of native ToHydA^ADT^, any catalytic activity in ToHydA^PDT^ is likely to fall below the detection limit. Experimental validation was therefore beyond the scope of this work, but it remains an interesting question for future studies.

## Conclusions

In conclusion, this study demonstrates the impact of ligand substitution at the [2Fe]_H_ site of ToHydA on protein structure and dynamics. Replacing the ADT ligand by PDT prevents the formation of the H_inact_ and H_trans_ states observed in ToHydA^ADT^. Our experimental and theoretical approach combining ATRFTIR spectroscopy and MD simulations highlights the role of the interaction between the ADT bridgehead and the C212 sidechain in the formation of the H_inact_ and H_trans_ states. In the absence of this interaction in ToHydA^PDT^ the thiol group of C212 is detached from Fe_d_, significantly reducing the population of the H_inact_ state. As-isolated ToHydA^PDT^ predominantly adopts the H_hyd_ state, which is atypical for this variant under such conditions. These findings provide valuable insights into ligand-dependent modulation of [FeFe]-hydrogenases, potentially opening the way towards applications in enzyme engineering and the development of biomimetic catalysts for hydrogen conversion.

## ASSOCIATED CONTENT

### Supporting Information

The supporting information includes experimental and simulation details, additional FTIR spectra of ToHydA^PDT^ and comparison of O_2_-treated spectra of ToHydA^ADT^ and ToHydA^PDT^.

## AUTHOR INFORMATION

### Author Contributions

The manuscript was written through contributions of all authors. All authors have given approval to the final version of the manuscript. All authors conducted research and/or analyzed data. S.G. expressed and isolated the proteins. S.G. and S.N. performed ATRFTIR spectroscopy. C.K.D. conducted all the MD simulations. S.G., C.K.D., S.N., L.V.S. and T.H. wrote and/or edited the manuscript.

### Notes

The authors declare no competing financial interest.

## ACKNOWLEDGMENT

S.G. thanks Deutscher Akademischer Austauschdienst (DAAD) for funding her doctoral scholarship. This project received funding from the Deutsche Forschungsgemeinschaft (DFG) under Germany’s Excellence Strategy – EXC 2033 – 390677874 – RESOLV. We thank Shanika Yadav and Ulf-Peter Apfel from the Faculty of Chemistry and Biochemistry at Ruhr University Bochum for their synthesis and provision of the [2Fe]^MIM^ (with both ADT and PDT ligands) used in in-vitro maturation.

