## Supporting Information for "How the Azadithiolate Ligand Impacts O_2_-Stability of Group B [FeFe]-Hydrogenase ToHydA"

### Experimental details

#### Expression and purification

The *E. coli* BL21 (DE3) ΔiscR[1] strain was transformed with a pET21b expression plasmid carrying a codon-optimized ToHydA gene to produce the apo form of ToHydA, which lacks the $[2Fe]_H$ subcluster.[2] Expression and purification were performed under strictly anaerobic conditions in the absence of the hydrogenase maturases HydE, HydF, and HydG, following previously established protocols.[2,3] The protein was purified using Strep-Tactin high-capacity resin (IBA GmBH) via affinity chromatography. Protein concentration was determined using the Bradford assay,[4] and its purity was assessed through SDS-PAGE.[5] The purified protein was stored at −80 °C in a 100 mM Tris-HCl buffer at pH 8 supplemented with 2 mM sodium dithionite (NaDT).

#### *In-vitro* maturation

For reconstitution, the ToHydA apoprotein was incubated on ice for 1 hour with a 10-fold molar excess of the synthetically prepared $[2Fe]^{MIM}$, the cofactor mimic of $[2Fe]_H$. Specifically, ToHydA was reconstituted with either the $[2Fe]^{MIM}$ with azadithiolate ligand, $[Fe_2[\mu\text{-}(SCH_2)_2NH](CN)_2(CO)_4]^{2-}$,[6] to form $ToHydA^{ADT}$, or the $[2Fe]^{MIM}$ with propanedithiolate ligand, $[Fe_2[\mu\text{-}(SCH_2)_2CH_2](CN)_2(CO)_4]^{2-}$,[7] to form $ToHydA^{PDT}$. The incubation was carried out in 100 mM potassium phosphate buffer ($K_2HPO_4/KH_2PO_4$, pH 6.8) supplemented with 2 mM sodium dithionite (NaDT), following an established protocol.[7] After incubation, the reconstituted holoenzymes were purified via size-exclusion chromatography using a NAP-5 column (GE Healthcare) to remove excess $[2Fe]^{MIM}$. The matured holoenzymes were then stored at −80 °C in 100 mM Tris-HCl buffer (pH 8) containing 2 mM NaDT.

#### ATR−FTIR spectroscopy

Attenuated total reflectance Fourier transform infrared (ATR-FTIR) spectroscopy was performed using a Bruker Tensor II spectrometer (Bruker Optik, Germany) equipped with a 9-reflection ZnSe/Si crystal (Microm ATR Vision, Czitek). All measurements were conducted under strictly anaerobic conditions (2% $H_2$, 98% $N_2$) at 25°C. FTIR spectra were recorded within a spectral range of 4000 to 1000 $cm^{-1}$ with a resolution of 2 $cm^{-1}$. For sample preparation, 4 µL of protein at a concentration of 0.5 mM was applied to the ATR crystal and allowed to dry under anaerobic conditions until the characteristic H-cluster absorption bands were observed (the resulting spectra referred to as as-isolated state of the protein). To maintain hydration and prevent excessive drying, a humidified gas stream containing $N_2$ (8 L/min) and $O_2$ (air, 2 L/min) was applied for approximately 10 minutes. For chemical treatments, 4 µL of 25 mM NaDT, prepared in 100 mM Tris-HCl buffer ($H_2O/D_2O$, pH 8), was added to the dried protein film. For primary isotope effect experiments, H→D exchange was achieved by replacing the standard buffer (100 mM Tris-HCl in $H_2O$ at pH 8) with deuterated buffer (100 mM Tris-HCl in $D_2O$ at pH 8). From the maturation step onward, $D_2O$-containing buffer was used in the NAP-5 column during buffer exchange to ensure complete H→D substitution while also removing excess $[2Fe]^{MIM}$. The as-isolated proteins consistently contained 2 mM NaDT, including those in $D_2O$-based buffer.

#### Molecular dynamics simulation details

All molecular dynamics (MD) simulations were performed using the GROMACS software package (version 2021.1).[8] The initial structure of the ToHydA WT [FeFe]-hydrogenase was obtained from previously published work. To generate the ToHydA$^{PDT}$ variant, the ADT ligand in the WT structure was replaced with a PDT ligand.

The simulations employed the CHARMM36 protein force field along with the CHARMM-specific TIP3P water model.[9] This study specifically examined the $H_{ox}$ state of ToHydA [FeFe]-hydrogenase. Force field parameters for the H-cluster and accessory FeS clusters, including coordinating cysteine and histidine residues, were derived from the work of Chang et al.,[10] with further refinements based on the recommendations of McCullagh and Voth.[11] The force field parameters for the ADT ligand were modified accordingly to accommodate the PDT ligand. Each protein was fully solvated in a system containing 16,519 water molecules, with charge neutrality maintained by the addition of 17 $Na^+$ ions. The simulations were conducted using periodic boundary conditions within dodecahedral simulation boxes. For the ToHydA$^{PDT}$ variant, the total system comprised 56,209 atoms.

The molecular dynamics (MD) protocols used in this study followed those employed in previous simulations. Prior to running MD simulations, the systems underwent energy minimization using 10,000 steps of the steepest descent algorithm. This was followed by a stepwise equilibration process with harmonic position restraints applied to specific atom groups. The equilibration phase began with an NVT simulation, where the system temperature was gradually increased from 0 K to 300 K over 0.2 ns. During this phase, harmonic restraints with force constants of 1,000 kJ/mol/nm$^2$ were applied to all non-hydrogen atoms of the protein and FeS clusters. The equilibration process then continued for 2.5 ns under NPT ensemble conditions. In the first 0.5 ns of NPT equilibration, the same position restraints were maintained on all non-hydrogen atoms of the protein and FeS clusters. During the remaining 2.0 ns of NPT equilibration, restraints were applied exclusively to the protein backbone atoms, allowing the side chains, FeS clusters, and surrounding water molecules to adjust and relax. For production simulations, three independent 1,000 ns MD runs were conducted for each of the three protein systems under NPT conditions at 300 K. These simulations were initialized from the equilibrated structures, with different random seeds used to assign initial atomic velocities according to a Maxwell-Boltzmann distribution. For subsequent analyses, only the final 500 ns of each simulation were considered.

Throughout the MD simulations, the system temperature was maintained at 300 K using the velocity rescaling thermostat developed by Bussi et al.,[12] with a coupling time constant of 0.1 ps. To sustain a constant pressure of 1 bar, an isotropic weak coupling Berendsen barostat was employed, featuring a coupling time constant of 2 ps and a compressibility of $4.5 \times 10^{-5}$ bar$^{-1}$.[13] Short-range Coulomb and Lennard-Jones (6,12) interactions were computed using a buffered Verlet pair list,[14] where interaction potentials smoothly transitioned to zero at a cutoff distance of 1.2 nm, with forces gradually shifted to zero between 1.0 and 1.2 nm. Longrange electrostatic interactions were handled using the particle mesh Ewald (PME) method, utilizing a grid spacing of 0.12 nm.[15] To enforce bond constraints, the LINCS algorithm was applied to all protein bonds involving hydrogen atoms,[16] while the SETTLE algorithm was used to constrain internal degrees of freedom in water molecules.[17] This setup enabled numerical integration of the equations of motion with a time step of 2 fs.

### Supporting figures and tables

Figure S1. FTIR spectra of as-isolated ToHydA$^{PDT}$ (replicate).

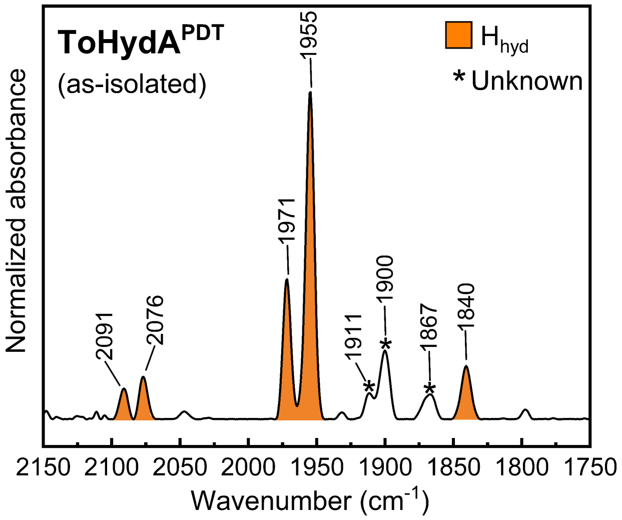

**Figure S1:** FTIR spectra of as-isolated ToHydA$^{PDT}$ (replicate). The spectra of the proteins are normalized to the second amide band (1535-1545 cm$^{-1}$). 0.4 - 0.5 mM protein sample was prepared in 100 mM Tris/HCl buffer at pH 8.0.

Figure S2. FTIR spectra of as-isolated, $H_2$-treated and 25mM NaDT-treated ToHydA$^{PDT}$ at pH 8 and pH 10.

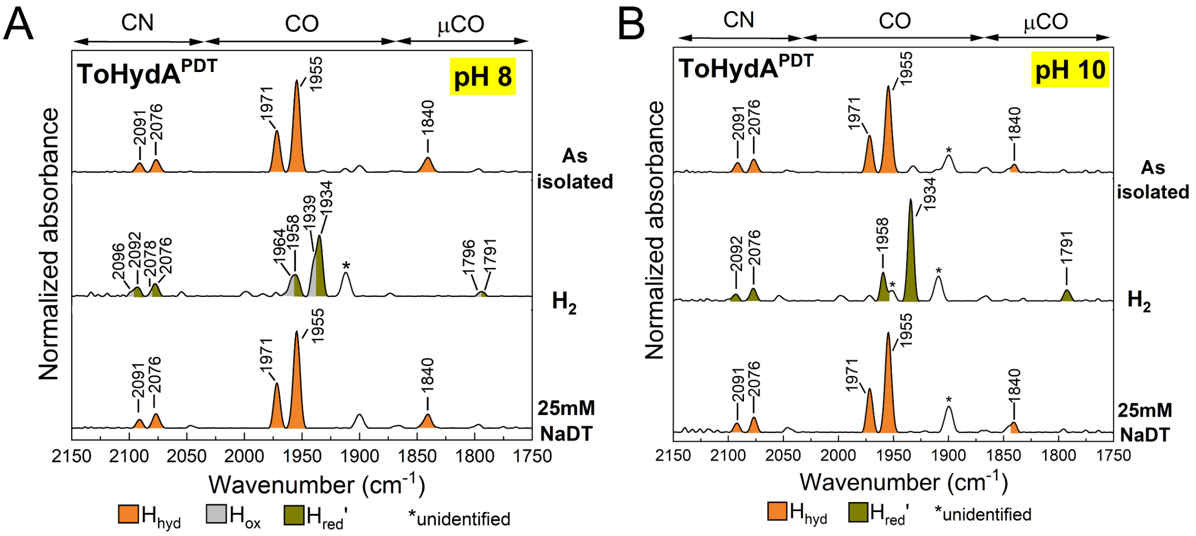

**Figure S2:** FTIR spectra of ToHydA$^{PDT}$ A) at pH 8 and B) at pH 10 in as-isolated state (upper panels), upon $H_2$ (5 L/min) purging (middle panels), and after 25 mM NaDT treatment (lower panels). The spectra of the proteins were normalized to the second amide band (1535-1545 cm$^{-1}$). 0.4 - 0.5 mM protein samples were prepared in 100 mM Tris/HCl buffers containing 2mM NaDT.

Figure S3. Overlay of FTIR spectra of ToHydA$^{ADT}$ and ToHydA$^{PDT}$ following $O_2$ treatment.

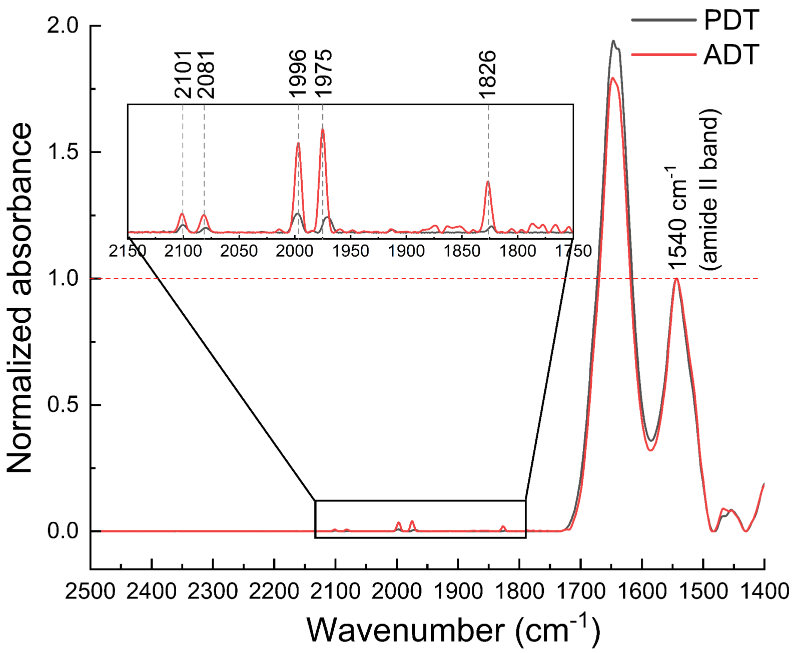

**Figure S3:** Overlapped spectra of ToHydA$^{ADT}$ and ToHydA$^{PDT}$ variant after $O_2$ (air, 2 L/min) exposure, spanning 2500–1400 cm$^{-1}$, both spectra normalized to the second amide band (1535–1545 cm$^{-1}$), indicated by a red horizontal dotted line. An enlarged view of the H-cluster region is shown in the rectangular inset. Peaks corresponding to the $H_{inact}$ state are marked by gray dotted lines and labeled accordingly.
